# Host factors influence the sex of nematodes parasitizing roots of *Arabidopsis thaliana*

**DOI:** 10.1101/273391

**Authors:** Muhammad Shahzad Anjam, Syed Jehangir Shah, Christiane Matera, Elżbieta Różańska, Miroslaw Sobczak, Shahid Siddique, Florian M.W. Grundler

## Abstract

Plant-parasitic cyst nematodes induce hypermetabolic syncytial nurse cells in the roots of their host plants. Syncytia are their only food source. Cyst nematodes are sexually dimorphic, with their differentiation into male or female strongly influenced by host environmental conditions. Under favorable conditions with plenty of nutrients, more females develop, whereas mainly male nematodes develop under adverse conditions such as in resistant plants. Here, we developed and validated a method to predict the sex of beet cyst nematode (*Heterodera schachtii*) during the early stages of its parasitism in the host plant *Arabidopsis thaliana*. We collected root segments containing male-associated syncytia (MAS) or female-associated syncytia (FAS), isolated syncytial cells by laser microdissection, and performed a comparative transcriptome analysis. Genes belonging to categories of defense, nutrient deficiency, and nutrient starvation were overrepresented in MAS as compared to FAS. Conversely, gene categories related to metabolism, modification, and biosynthesis of cell walls were overrepresented in FAS. We used β-glucuronidase (GUS) analysis, qRT-PCR, and loss-of-function mutants to characterize FAS- and MAS-specific candidate genes. Our results demonstrated that various plant-based factors, including immune response, nutrient availability, and structural modifications, influence the sexual fate sex determination of cyst the nematodes.

## Introduction

Cyst nematodes are plant parasites that induce specialized syncytial feeding structures inside the roots of their host plants. Syncytia are the only food source for developing juveniles and adult females. Cyst nematodes are sexually dimorphic with mobile males and sedentary females. Under field conditions the proportion of male and female nematodes is of fundamental importance. A high number of females usually lead to increased crop damage and heavy soil infestation. High numbers of males usually occur on resistant plants and indicate adverse conditions for the nematodes. The mechanism of sex determination in this group of plant parasitic nematodes is, however, not clearly understood. Therefore, a better understanding of the factors behind sexual differentiation could be helpful in developing methods to exploit plant factors that lead to male formation.

The infective second-stage juveniles of cyst nematodes (J2) invade the roots in the elongation zone behind the root tip. Once inside the root, stylet thrusts and the release of cell wall degrading enzymes and other proteins from the nematode’s pharyngeal secretory glands facilitate its intracellular movement through different tissue layers of the host, towards the vascular cylinder. Upon reaching the vascular cylinder, the nematode probes single cells to select a suitable initial syncytial cell (ISC) (Golinowski, Grundler & Sobczak, 1996; Sobczak, Golinowski & Grundler, 1999). Once the ISC is selected, the nematode becomes sedentary and a cocktail of secretions is released into the ISC that manipulates plant defense and metabolic pathways leading to the development of syncytium. Subsequently, nematodes develop into males or females over the period of three molts (J3, J4, and adults). The adult male nematodes leave the roots, but the females remain sedentary and produce eggs after fertilization. Finally, the females die, and their bodies turn into egg-protecting cysts. Because of the prolonged sedentary phase required for reproduction, female cyst nematodes require, on average, 29 times more food compared to males (Müller, Rehbock, & Wyss, 1981). Further, female-associated syncytia (FAS) are larger than male-associated syncytia (MAS).

A considerable amount of work has been carried out to clarify the influence of environmental and genetic factors on the sex of cyst nematodes. The results, however, are not all in agreement. The first report on variations in the sex ratio was provided by Molz (1920), who found that the sex of the sugar beet cyst nematode *Heterodera schachtii* is strongly influenced by the physiological state of the host plants, leading to the suggestion of environmental sex determination (ESD) in cyst nematodes. However, contrasting conclusions were drawn a few years later by Sengbusch (1927), who repeated some of Molz’s experiments and suggested that the high percentages of males under unfavorable conditions result from a differential death rate of the female larvae. This conclusion was based on his estimations that females require 35 times more food as compared to males. Later, Ellenby (1954) published the results of experiments with *H. rostochiensis* (syn. *Globodera rostochiensis*) on potato roots and argued that if a variation in the sex ratio is due to the differential death rate of females, then the bodies of the deceased females should be present in the roots. Accordingly, he designated all nematodes that were not adult males as females. Under this determination, the proportion of males to females increased strongly with an increased intensity of unfavorable environmental conditions, thus reinforcing the ESD view. Adding a single juvenile of *G. rostochiensis* to the host root resulted in an overwhelming majority of females in two independent studies, which was attributed to a decrease in competition for feeding site induction (Den Ouden, 1960; Trudgill, 1967). Grundler, Betka, & Wyss (1991) performed a number of experiments by adding a single juvenile of *H. schachtii* on *Brassica rapa* growing in a nutrient solution containing minerals and various concentrations of sucrose. They also found that the majority of juveniles developed into females under favorable conditions. Although single-juvenile experiments indicated that ESD plays a role in the sexual outcome of cyst nematodes, in the majority of these experiments, the number of males remained rather constant, whereas the number of females fluctuated under different environmental and nutritional conditions.

Transcriptome analysis is often the first step towards identifying genes and pathways that underlie a biological phenomenon. A transcriptome and proteome analysis of FAS induced by *H. schachtii* in *Arabidopsis* roots provided valuable insights into the molecular functioning of syncytium (Szakasits et al., 2009; Hütten et al., 2015). However, it is technically challenging to compare the differences between MAS and FAS with this approach. For one thing, when the morphological features that allow for sex determination become apparent, the nematodes are already sexually differentiated into males and females (Raski, 1950; Wyss, 1992); therefore, it is unlikely that a transcriptome analysis at this time point would provide insights into the host factors that influence sexual differentiation. In addition, the isolation of pure syncytial material during the early stages of syncytium development is challenging and laborious.

Here, we established and validated a strategy to predict the sex of *H. schachtii* juveniles during the early stages of infection when their sexual outcome is not yet apparent (Wyss, 1992). We isolated pure syncytial material via laser capture microdissection (LCM), which allowed us to compare the transcriptomes of potential MAS and FAS and identify sets of genes that are differentially regulated during the early stages of syncytium development. Our subsequent infection assays of knock-out mutants corresponding to the differentially-regulated genes provided new insights into plant-based factors that influence sexual differentiation of *H. schachtii*.

## Materials and Methods

### Plant growth conditions

*Arabidopsis thaliana* seeds were surface sterilized using 0.7% NaOCl (v/v) for 5 minutes followed by three successive washings with sterile water. The sterilized seeds were grown in Petri dishes containing Knop medium supplemented with 2% (w/v) sucrose (Sijmons, Grundler, von Mende, Burrows, & Wyss 1991). The plates were incubated in growth room at 25 °C, with an alternating period of sixteen hours of light and eight hours of dark under sterile conditions.

### Nematode measurement, prediction and infection assay

10-days-old plants were infected with freshly hatched J2s of *Heterodera schachtii*. For relative growth, the J2s that invaded and established in the lateral roots at 24 h after inoculation (hai) were marked with permanent markers on Petri dishes and imaged daily for the next five days. Their feeding establishment was defined when a nematode stopped stylet movements. The average size of a nematode was measured as previously described (Siddique et al., 2014). For each experiment, 40 - 50 nematodes were measured and the experiments were repeated three times. An empirical curve was drawn across average measurements and the sex of the nematodes was re-evaluated at 12 days post infection (dpi). Similarly, for prediction assays J2s established feeding sites in lateral roots were marked at 24 hai and observing their relative development were predicted at 4 and 5 dpi. The only juveniles associated with single syncytium were considered for prediction. The sex of predicted juveniles was confirmed at 12 dpi. For infection assays, the numbers of males and females were counted. Moreover, the average size of females and the average size of syncytia at 14 dpi were measured as described recently (Anwer et al., 2017).

### Sample collection, processing and microarray data analysis

Root segments containing putative MAS or FAS were collected in a Farmer’s fixative solution on ice. The samples were embedded in an optimum cutting medium (OCT) (Polyfreez^®^) using Tissue´Tek^®^ cryo molds and sections of 10 µm were cut and the total RNA was extracted as described previously (Anjam et al., 2016). A cDNA synthesis was performed with NuGEN’s Applause 3’-Amp System (Cat. No. 5100), according to the manufacturer’s instructions, and started with 100 ng of total RNA. NuGEN’s Encore Biotin Module (Cat. No. 4200-12) was used to fragment 3.95 µg cDNA followed by Biotin-labelling according to the manufacturer’s instructions. Hybridization, washing, and scanning, were performed according to the Affymetrix 30 GeneChip Expression Analysis Technical Manual (Affymetrix, Santa Clara, CA, USA). Three chips were hybridized for control and infected samples, with each microarray representing an independent, biological replicate. Primary data analysis was performed with Affymetrix software Expression Console v1.* using the MAS5 algorithm. Statistical analysis of microarray data was performed as described previously (Mendy et al., 2017). The gene set enrichment analysis (GSEA) of FAS and MAS data was performed using PlantGSEA tool kit as described previously (Yi, Du, & Su, 2013). The genes with fold change >1.5 and P-value <0.05 were selected for enrichment analysis in GO (gene ontology) annotations of domain ‘biological process’. We compared expected and actual percentage of gene enrichment in selected subcategories. The number of *Arabidopsis* genes used for calculating expected enrichment was 27029. GSEA used these genes from TAIR database annotated into 7041 GO gene sets (Yi et al., 2013). The actual enrichment was calculated from 124 FAS and 331 MAS genes. The expression of top 100 FAS and MAS genes in different anatomical parts, under nutrient and biotic stress conditions, was analyzed by genevestigator.

### Genotyping and expression analysis of knock-out mutants

Single T-DNA inserted knock-out mutants for selected genes (Supplementary Table S1) were ordered from the relevant stock center. The homozygosity of SALK mutant lines (NASC, The European Arabidopsis Stock Centre, www.arabidopsis.info) was confirmed via PCR using primers given in Table S2. The homozygous mutants of SALK and GABI-KAT lines (University of Bielefeld, Germany, www.gabi-kat.de) were confirmed to be completely absent for expression of required gene through RT-PCR with gene specific primers given in Table S2.

### Development of promoter-reporter lines and GUS analysis

Promoter regions upstream of the 5’ UTR of *LPTG-6* (1361 bp), *BGLU28* (1471 bp) and *CWLP-1* (1214 bp) were amplified by Gateway PCR using *Arabidopsis* Col-0 genomic DNA as template using primer given in Supplementary Table S3. Subsequently, promoters were cloned via gateway cloning upstream of *GUS* gene in pMDC162. *Promoter::GUS* constructs were introduced into *Agrobacterium tumefaciens* GV3101 for transformation of *Arabidopsis* Col-0 plants by the floral dip method. T3 homozygous lines were generated and analyzed for GUS expression as described recently (Shah et al., 2017). Four plants from each of three transgenic lines per construct were analyzed at each time-point.

### Real time PCR

RNA from LCM-derived MAS or FAS was isolated and cDNA was amplified. Transcriptome abundance for candidate genes was analyzed using StepOne Plus Real-Time PCR System (Applied Biosystems, USA) as described recently (Anwer et al., 2017). β-Tubulin and ubiquitin were used as internal controls. Relative expression was calculated by the Pfaffl’s method (Pfaffl, 2001), where the expression of the candidate gene was normalized to the internal control to calculate fold change. Primer sequences for all genes are provided in Supplementary Table S4.

### Anatomy and ultrastructure

*Arabidopsis* Col-0 plants were grown and inoculated aseptically with freshly hatched J2s of *H. schachtii* as described above. Root segments containing putative MAS or FAS were dissected at 5 dpi, processed for light and transmission electron microscopy, and examined as described previously (Golinowski et al., 1996).

## Results

### J2 females grow faster than J2 males

To determine whether there is a difference in growth patterns between male and female J2s, we grew plants *in vitro* and inoculated them with nematodes. We marked the juveniles that had successfully established the ISC at 24 hai and monitored their growth daily over the following 5 days. We were able to determine the sex of the marked nematodes at 12 dpi, and calculated the growth curves for males and females. Until 3 dpi, there was no difference in the average size between male and female juveniles; however, the female juveniles were significantly larger from 4 dpi and grew faster than male juveniles (Figure 1). These observations suggested that the sexual fate of the J2s may be determined during the first 4 – 5 days after infection. These results also suggested that it might be possible to predict the sex of a juvenile during the early stages of infection based on the differences in their sizes.

**Figure 1:**
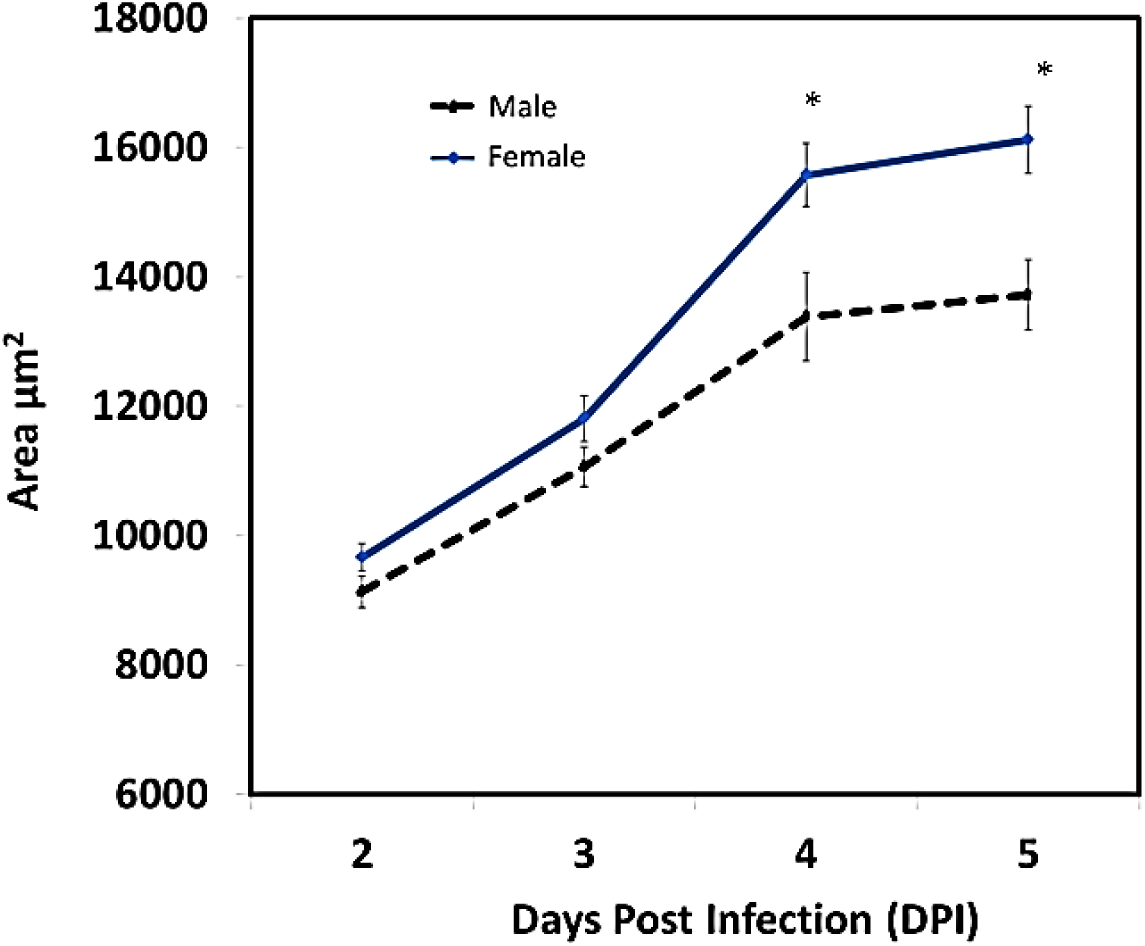
J2 females grow faster than J2 males. The nematodes that established an ISC in roots were marked and their growth was monitored over the next five days. The values represent the average size of a nematode +/− SE (n = 3). Data were analysed using two-tailed *t-test* (p < 0.05). Asterisks represent statistically significant differences compared to the control.

### The sex of nematodes can be accurately predicted as early as 5 dpi

To assess whether we can predict the sex of J2 nematodes during the early stages of infection, we inoculated the plants with nematodes and marked successfully established nematodes at 24 hai. The development of nematodes was assessed at 4 dpi and the sex of the nematodes was predicted as male or female at 5 dpi. The predicted sex of the juveniles was re-evaluated at 12 dpi, a time point where morphological differences between males and females can be clearly differentiated. We were able to predict the sex of female nematodes at 5 dpi with more than 90% accuracy, while the sex of male nematodes was correctly predicted with approximately 85% accuracy. Thus, the sex of the juveniles can be predicted successfully during the initial stage of infection (Table 1 and Figure 2).

**Table 1:** Sex prediction assays for *H. schachtii*. The experiment was repeated thrice independently and percentage of right prediction was calculated. +/− indicates standard deviation.

| Nematode | Prediction | Actual | Success (%) |
| --- | --- | --- | --- |
| Male | 76 | 66 | 86.3 ± 1.8 |
| Female | 125 | 115 | 91.2 ± 1.7 |

**Figure 2:**
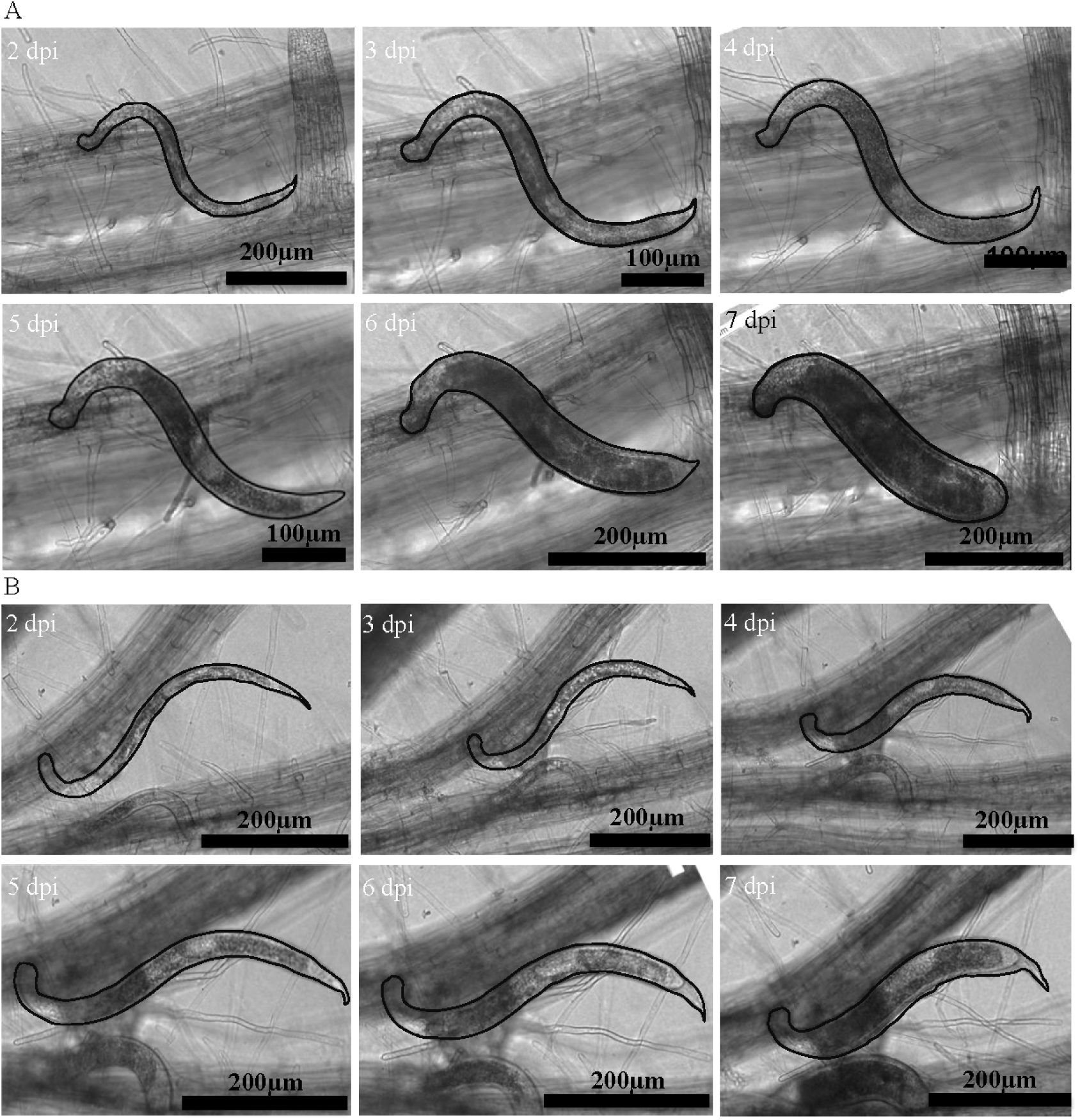
The sex of nematodes can be accurately predicted as early as 5 dpi. (A) Development of a female juvenile from 2 to 7 dpi. (B) Development of a male juvenile from 2 to 7 dpi. Note: a line has been drawn around the highlighted nematode to make stand out from other nematodes in the background. Scale bars = 200 µm.

### Structural and cellular differences between FAS and MAS

To investigate the cellular and ultrastructural differences between MAS and FAS, we predicted the sex of the nematodes at 5 dpi and dissected root segments containing syncytium and attached nematodes. The root segments were serially cross-sectioned and analyzed through light and transmission electron microscopy. We found that both MAS and FAS expanded via incorporation of hypertrophied vascular cylinder cells, but they differed strongly in anatomy (Figure 3A–3J). MAS were shorter and composed of fewer cells (Figure 3A–3E) than FAS (Figure 3F–3J). At the leading edge of the syncytium, where new cells were incorporated into axially expanding syncytium, recently incorporated cells were only slightly hypertrophied in MAS (Figure 3A), whereas their hypertrophy was more pronounced in FAS (Figure 3F). This difference was also clearly recognizable in submedian and median parts of both syncytia (Figure 3B and 3C versus Figure 3G and 3H). In the region close to the tip of the nematode’s head where the ISC had been selected, MAS usually had a crescent-like shape (Figure 3D) and many degraded cells were present. By contrast, the FAS were centrally located inside the vascular cylinder (Figure 3I). Below the nematode head, in the juvenile migration region, the extent of damaged cells was very high around predicted male juveniles (Figure 3E), whereas the vascular cylinder was more intact near the predicted female juveniles (Figure 3J). Another anatomical feature differentiating MAS and FAS was the very weak development of periderm-like secondary cover tissue around MAS (Figure 3A–3C versus Figure 3F–3I), which consisted of 3–4 continuous cell layers sheathing the FAS (Figure 3F–3H), but only 1–2 cell layers in MAS (Figure 3A-3C).

**Figure 3:**
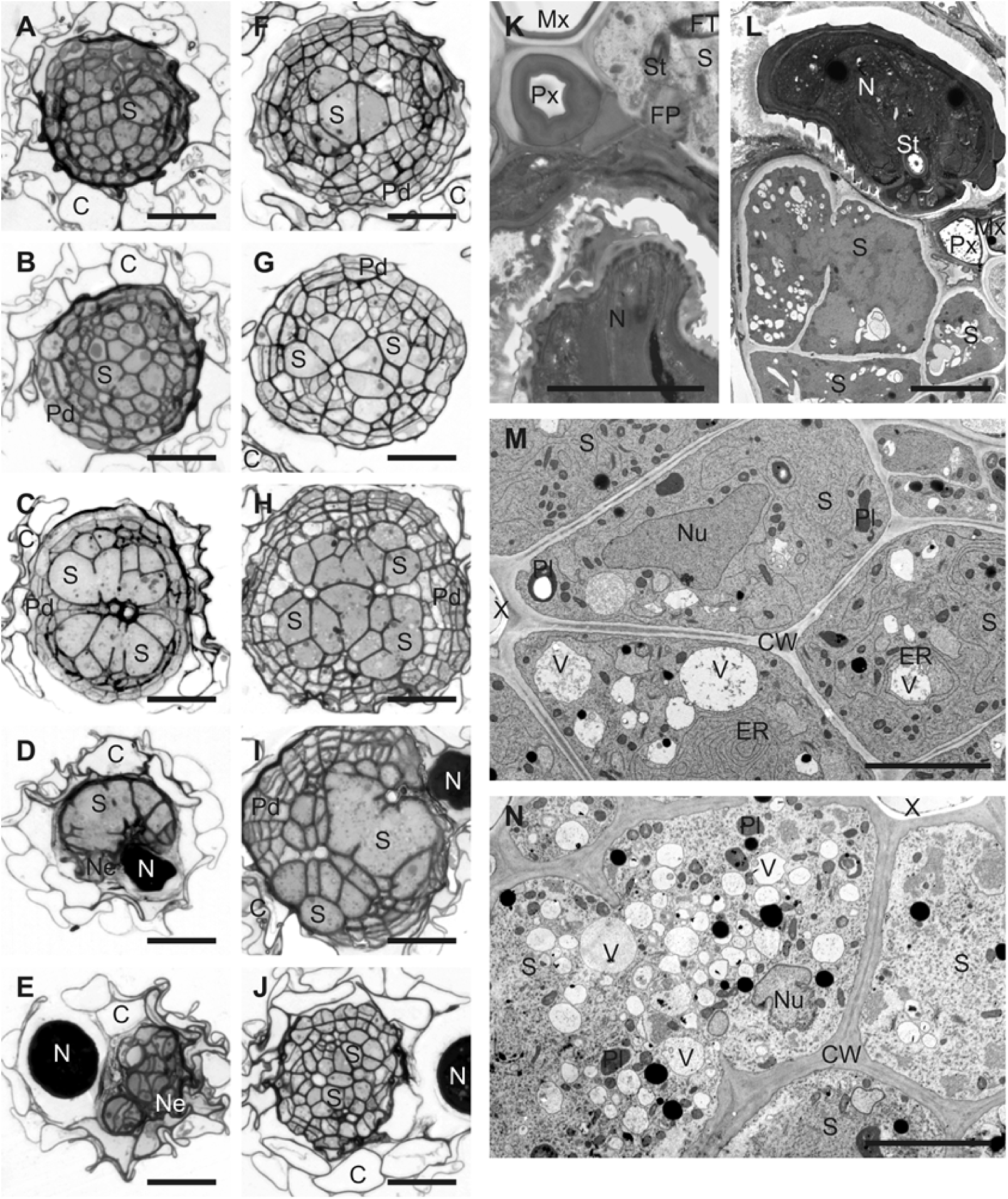
Structural and cellular differences are observed between FAS and MAS. Light (A–J) and transmission electron microscopy (K–N) images taken from cross sections of syncytia associated with male (A–E, L, and N) and female (F–J, K, and M) juveniles. The images were taken from sections made at the leading edge of syncytium (A and F), the submedian region of syncytium (B and G), the median region of syncytium (C, H, M, and N), next to the nematode’s head (D, I, K, and L), and along the nematode migration path (E and J). Abbreviations: C-cortex, CW-cell wall, ER-cistern of endoplasmic reticulum, FP-feeding plug, FT-feeding tube, Mx-metaxylem vessel, N-nematode, Ne-necrosis, Nu-nucleus, Pd-periderm-like tissue, Pl-plastid, Px-protoxylem vessel, S-syncytium, St-stylet, X-xylem, V-vacuole. Scale bars = 50 µm (A–J) and 5 µm (K–N).

Serial sectioning of syncytia allowed us to localize the heads and stylets of several predicted male and female juveniles. The identification of ISC at this time point (5 dpi) was difficult due to extensive cell enlargement and cell wall openings formation. However, juveniles appeared to always select their ISCs in cells next to the xylem vessels (Figure 3I, 3K and 3L). FAS were induced among procambial cells (Figure 3I and 3K), whereas MAS were induced among pericyclic cells (Figure 3L).

Ultrastructural analysis showed that at 5 dpi, FAS had electron-dense cytoplasm with notably smaller vacuoles (Figure 3M). The protoplasts of FAS contained enlarged and amoeboid nuclei, and numerous organelles. By contrast, MAS had less electron-dense cytoplasm almost completely devoid of rough ER cisternae, and small vacuoles were more numerous than in FAS (Figure 3N).

### FAS and MAS have transcriptional differences

To reveal changes in gene expression between FAS and MAS during the early stages of infection, we performed GeneChip analyses. Plants were grown and inoculated with nematodes. Invading J2s that had successfully established the ISC (as defined by a cessation of stylet movements) were marked at 24 hpi, and the sex was predicted at 4 dpi. Root segments containing potential MAS or FAS were dissected at 5 dpi. The samples were embedded in an optimum cutting medium and pure syncytial material was isolated using laser capture microdissection (LCM). Total RNA was extracted, labelled, amplified, and hybridized with the *Arabidopsis* ATH1 GeneChip designed for the detection of 24,000 genes. We compared the transcriptomes of FAS and MAS and found that 455 genes were differentially expressed (fold change >1.5), with a false discovery rate below 5% (Table S5). A higher number of genes showed increased transcript abundance in MAS (331), as compared to FAS (124). FAS- or MAS-specific are used throughout this manuscript to describe genes that have an increased expression in FAS or MAS. Lists of the 50 most strongly differentially-expressed genes are given in Tables S6 and S7.

### Defense- and nutrient stress-related genes are overrepresented in MAS

To understand the processes that are altered between FAS and MAS, we performed a GO enrichment analysis of FAS- or MAS-specific genes by computing overlaps with 7,041 previously defined data sets using PlantGSEA software (Yi et al., 2013). Categories that were particularly enriched in the genes upregulated in FAS include: ‘polysaccharide biosynthetic processes’, ‘carbohydrate biosynthetic and metabolic processes’, ‘polysaccharide metabolic processes’, and many ‘cell wall-related biosynthetic and metabolic process’ categories (Figure 4A). For the MAS-specific genes, categories of ‘response to stimulus’, ‘response to stresses’, ‘response to starvation’, ‘response to nutrition’, and ‘defense and innate immune response’, were significantly overrepresented (Figure 4B). The complete list of overrepresented categories in GO enrichment for FAS and MAS is given in Supplementary Tables S8 and S9.

**Figure 4:**
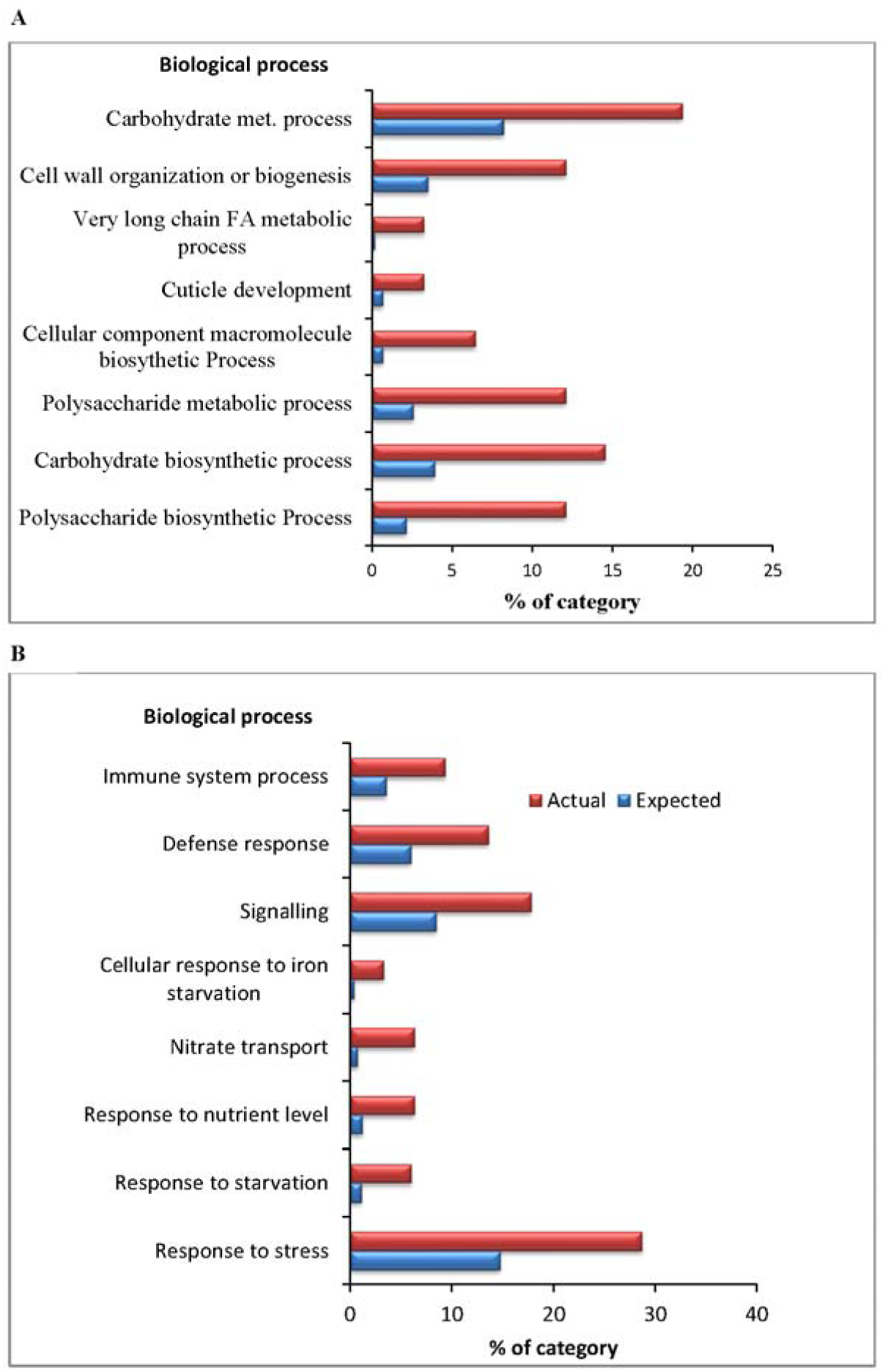
GO enrichment analysis showed overrepresentation of defense- and nutrient stress-related genes in MAS. The percentage of expected and actual genes found in the examined subset is shown on the x-axis. The blue bar represents the expected set of genes matching the respective category and red bar indicates the set of differentially regulated genes in the current study. All gene sets shown have a FDR < 0.05 and a fold change < 1.5. (A) Overrepresentation of FAS-preferentially upregulated genes (124) with respect to the Arabidopsis genome. (B) Overrepresentation of MAS-preferentially upregulated genes (331) with respect to the Arabidopsis genome.

Next, we used Genevestigator (Zimmerman et al., 2014) to analyze the expression of FAS-specific and MAS-specific genes in response to biotic infections and found that many of the MAS-specific genes were strongly induced in response to biotic infections (Supplementary Figure S1), while FAS-specific genes showed little to no induction in response to biotic infections (Supplementary Figure S2). We performed a similar analysis to evaluate the expression patterns of FAS-specific or MAS-specific genes in response to nutrient stress. Whereas the FAS-specific genes were not related or only slightly related to nutrient stress responses (Supplementary Figure S3), the MAS-specific genes showed a clear tendency to be induced by iron or sulfur deficiency. Interestingly, the expression of many of the MAS-specific genes was suppressed when seedlings were treated with sucrose or glucose (Supplementary Figure S4).

### Validation of GeneChip data by quantitative RT-PCR

The expression of eight FAS-specific or MAS-specific genes was further validated using quantitative RT-PCR. Our qRT-PCR analysis showed the same trend as that indicated by our microarray analysis (Table 2). However, the fold change for some candidate genes (*CWLP1, CELL WALL PLASMA MEMBRANE LINKER PROTEIN*; *GH3.3, INDOLE-3-ACETIC ACID-AMIDO SYNTHETASE*; *PGIP1, POLYGALACTURONASE INHIBITOR PROTEIN 1*; and *LNG1, LONGEFOLIA 1*) was much higher as compared to the microarray results.

**Table 2:** Validation of microarray results by qRT-PCR. The fold change in expression of candidate genes in FAS when compared to MAS. The values represent average for three biological replicates +/− S.E.

| Gene Name | Fold Change (Female vs Male) |  |
| --- | --- | --- |
|  | Microarrays | qRT-PCR |
| BGLU-28 | 0.051 | 0.09 +/- 0.045 |
| GH3.3 | 3.92 | 8.72 +/- 1.74 |
| CWLP-1 | 13.46 | 36.25 +/-12.25 |
| PGIP1 | 0.37 | 0.29 +/- 0.08 |
| PGIP2 | 0.54 | 0.38 +/- 0.13 |
| PDF1.4 | 0.23 | 0.77 +/- 0.15 |
| LTPG6 | 2.65 | 2.94 +/- 1.24 |
| LNG1 | 2.58 | 6.05 +/- 1.20 |

### *Promoter::GUS* analysis

We generated promoter::GUS lines to analyse the spatio-temporal expression of two highly expressed candidate genes, one each for FAS (*CWLP1*) and MAS [*B-GLUCOSIDASE 28* (*BGLU28*)]. One representative homozygous line for each gene was infected with nematodes. The sex of the nematodes was predicted at 5 dpi and tissues were stained for GUS activity at 5 and 12 dpi (Figure 5). In *pCWLP1::GUS* plants, the majority of the FAS showed moderate to high GUS staining at 5 and 12 dpi, while only weak staining was detected in MAS (Figure 5). For the *pBGLU28::GUS* plants, FAS showed weak GUS staining, whereas MAS were stained strongly. In the uninfected controls, *pCWLP1::GUS* plants had slight GUS staining in the vascular cylinder; however, faint GUS staining was detected in root tips of both *pCWLP1:GUS* and *pBGLU28:GUS* plants (Figure 5).

**Figure 5:**
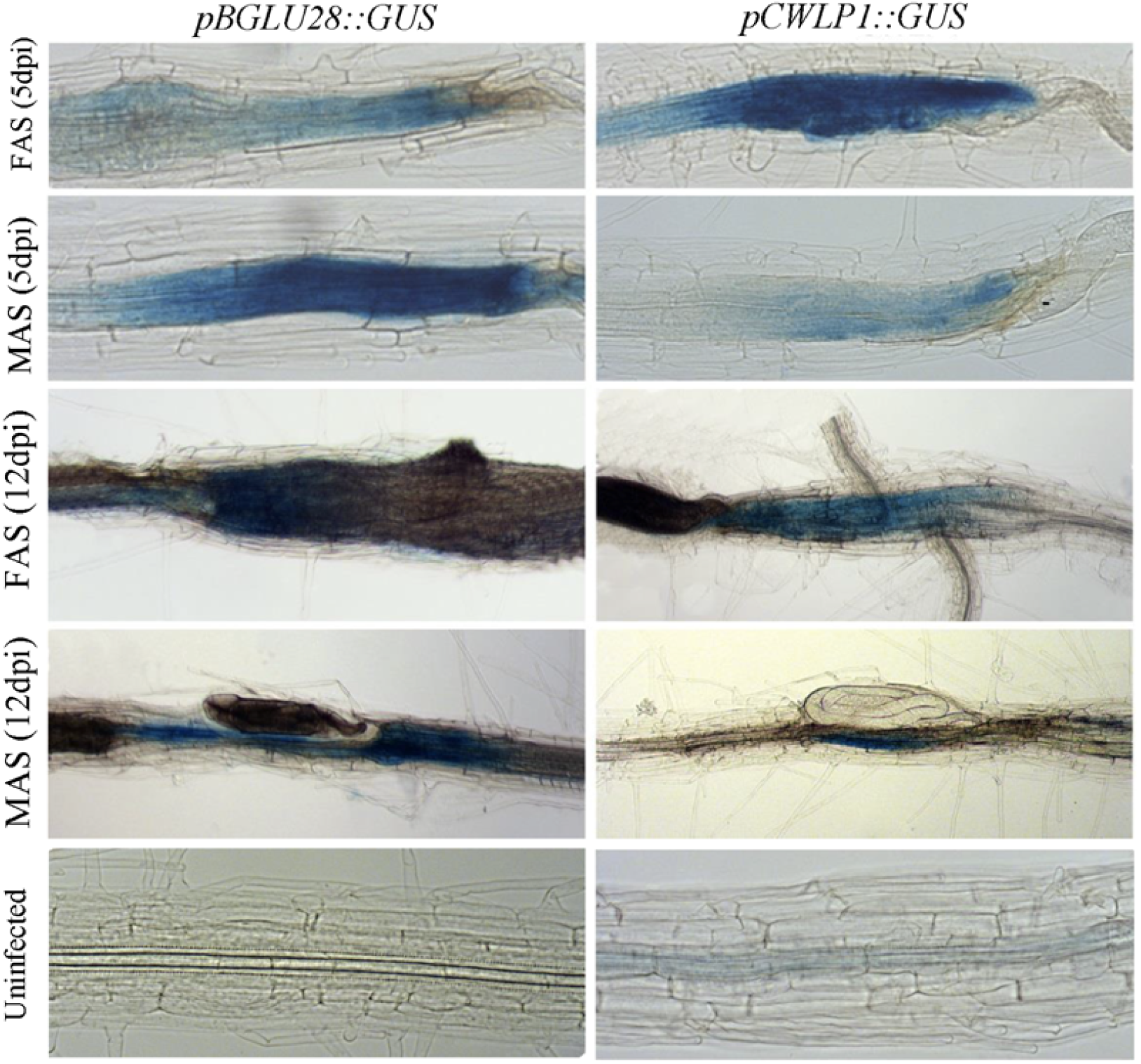
GUS staining of FAS and MAS at 5 and 12 dpi validates microarrays analysis. The figures on the left side represent the *pBGLU28::GUS* line and on the right side represent the *pCWLP1::GUS* line of *Arabidopsis*.

### Knocking out host candidate genes alters the sexual fate of nematodes

To investigate which genes/pathways are involved in ESD of nematodes, we selected seven FAS-specific genes [*CWLP1, MLP-like protein 423* (*MLP423*), *GLYCEROPHOSPHODIESTER PHOSPHODIESTERASE LIKE 5 (GDPDL5), LACCASE 11* (*LAC11*), *LNG1, IRREGULAR XYLEM 12 (IRX12)*, and *GLYCOSYLPHOSPHATIDYLINOSITOL-ANCHORED LIPID PROTEIN TRANSFER 6* (*LPTG6*)] and three MAS-specific genes [*BGLU28, BGLU30/DIN2*, and *BASIC HELIX-LOOP-HELIX TRANSCRIPTION FACTOR 101* (*BHLH101*)] for further characterization using loss-of-function T-DNA mutants (Supplementary Table S1 and Supplementary Figures S5). Plants were screened for the development of males and females at two weeks after inoculation. Out of the eight mutant lines for FAS-specific genes, two lines, *lng1* and *irx12*, showed a significant reduction in the number of females and a significant increase in the number of males as compared to the wild type Col-0. In addition, the *lptg6* mutant also showed a significant decrease in the number of females as compared to that of Col-0, but the number of males did not change significantly (Figure 6A). We also found that the size of the syncytia, but not the size of the females, was significantly reduced in the *ltpg6* and *lng1* mutants in comparison to Col-0 plants (Figure 6B–C). By contrast, no changes in nematode numbers were detected in the mutant lines for MAS-specific genes.

**Figure 6:**
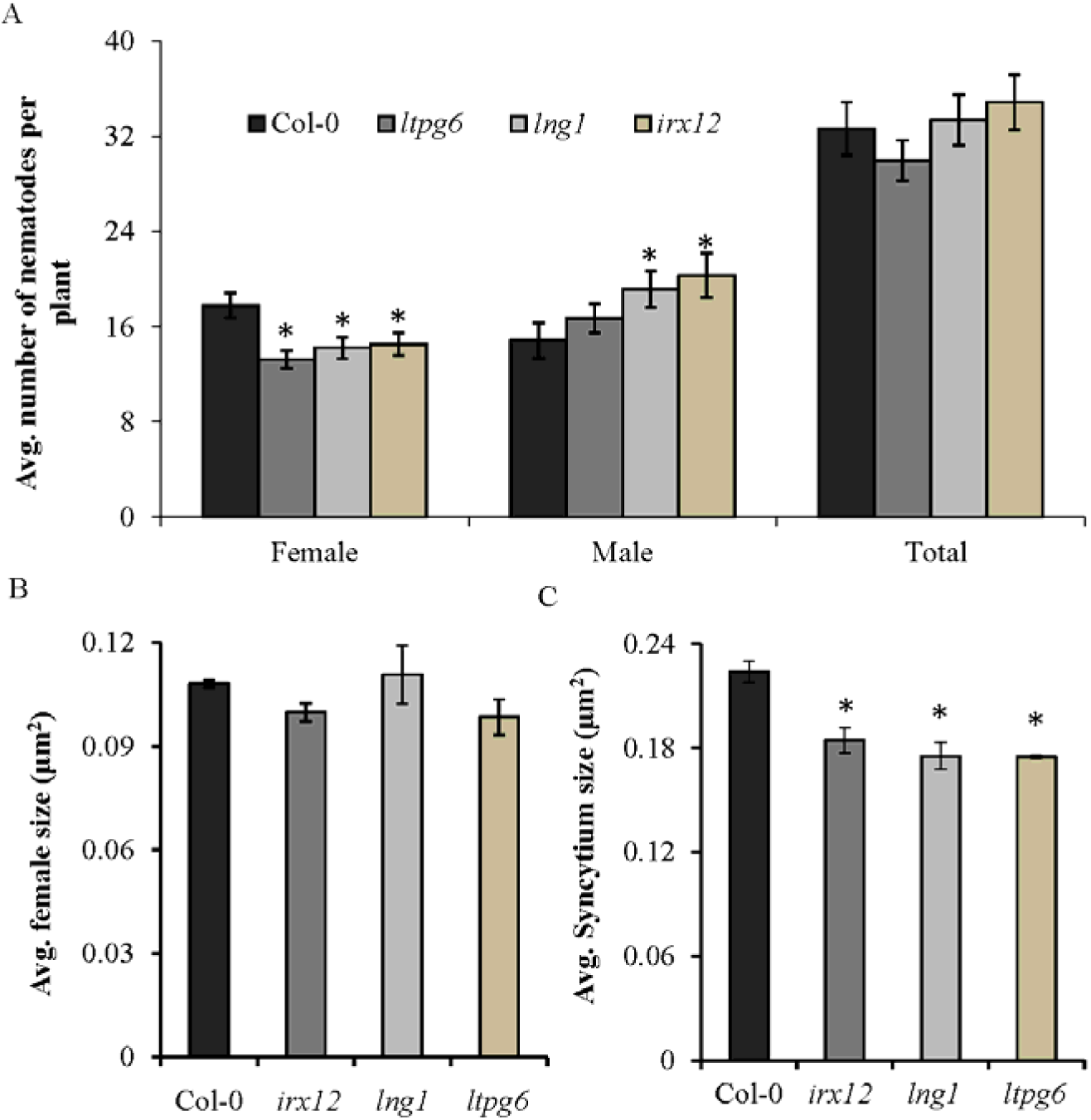
Knocking out host candidate genes alters the sexual fate of nematodes. (A) Average number of males, females, and total nematodes. (B) Average syncytium size. (C) Average female size. (A–C) Bars represent mean +/− SE from three independent biological replicates (n = 3). Data was analyzed using two-tailed *t-test*. Asterisks represent significant difference from Col-0 (at *P* < 0.05).

## Discussion

Here, we developed and validated a strategy to predict the sex of cyst nematodes with high certainty during the early stages of infection. The J2 nematodes that developed at the fastest rate during the first 4–5 days after syncytium induction became females, while those that grew slower became mainly males. Interestingly, a study by Müller et al. (1981) on comparative food consumption by male and female juveniles from roots of *B. napus* found that females consume about 29 times more food than males. Based on our data and previous literature, we concluded that the difference in food consumption a leads to the difference in body volume between the sexes.

### Anatomical and ultrastructural differences between FAS and MAS

Microscopic examination showed that FAS and MAS clearly differ in their anatomical organization. MAS were smaller in diameter and length in comparison to FAS. This corresponds to a smaller volume of the syncytium, and therefore to a smaller volume of protoplast on which the juveniles feed. Smaller syncytia also have a proportionally reduced surface area of the interface to the vascular cylinder’s conductive elements, and thus the influx of nutrients to MAS may also be reduced. Predicted male juveniles usually migrated inside the vascular cylinder for long distances and caused very extensive cell destruction, which raises the question of whether the continuity of xylem and phloem conductive elements was disrupted. If the conductive system was disrupted, the MAS would be located at its termini, which could affect the access of male nematodes to a continuous flow of nutrients. At the cellular level, the most obvious differences between MAS and FAS was the presence of numerous, small vacuoles in the former and cisternae of rough ER in the latter. In general, cisternal rough ER is implicated in the biosynthesis of proteins for secretion, whereas small vacuoles, in conjunction with a low electron density of syncytial cytoplasm, might indicate protoplast degradation related to a defense response or programmed cell death. These structural differences indicate that MAS have problems related to nutrient availability or defense responses resembling those observed in resistant plant–nematode interactions (Endo, 1965; Trudgill, 1991).

### Comparison of the transcriptomes of FAS and MAS

Earlier transcriptomic studies of syncytia induced in roots of soybean and Arabidopsis revealed that nematodes trigger massive changes in key pathways of the host transcriptome including pathways of hormonal regulation, cell wall architecture, cytoskeleton, and dedifferentiation of cells, facilitating the syncytium to function as metabolically highly active cells (Puthoff, Nettleton, Rodermel, & Baum, 2003; Ithal et al., 2007; Szakasits et al., 2009). However, these studies were conducted exclusively on FAS or a mixture of FAS and MAS. We hypothesized that there may be differences in transcriptional regulation between FAS and MAS that in turn influence the sexual differentiation of juveniles. Therefore, we performed a microarray analysis to compare the transcriptomes of MAS and FAS at 5 dpi.

### FAS undergoes much intense cell wall modifications as compared to MAS

Starting from a single cell, syncytia undergo extensive expansion via dissolution of the cell walls of neighboring cells. The outer wall of the syncytium is thickened to withstand increased turgor pressure inside the cell (Siddique, Sobczak, Tenhaken, Grundler, & Bohlmann, 2012, Böckenhoff & Grundler, 1994). Syncytial cell walls in contact with xylem vessels develop numerous ingrowths to enhance the surface area for absorption of nutrients and water (Golinowski et al., 1996; Offler, McCurdy, Patrick, & Talbot, 2003). These changes in cell wall structure help to meet the growing demand for food of developing nematodes. Microarray data published by Puthoff et al. (2003) at 3 dai and Szakasits at al. (2009) at 5 and 15 dpi also showed high upregulation of genes for ‘cell wall biosynthesis and modifications’. Our results indicate that FAS undergo a stronger upregulation of genes for ‘cell wall biosynthesis’, ‘metabolism’, and ‘modifications’ as compared to MAS. This is consistent with the finding that female nematodes require more food and grow much faster in comparison to males (Müller et al., 1981; Hofmann, Szakasits, Blöchl, & Sobczak, 2007), thus the upregulation of cell wall-related genes may support the higher nutritional requirement. This hypothesis was also supported by previous studies in which FAS contained more conspicuous cell wall ingrowths (transfer cells) and cell wall openings as compared to MAS (Golinowski et al., 1996; Sobczak, Golinowski, & Grundler, 1997).

### Immune responses are activated in MAS as compared to FAS

Invasion of the root by nematodes, and subsequent migration towards the vascular cylinder, cause cellular damage and activate plant defense responses (Holbein, Grundler, & Siddique, 2016; Mendy et al., 2017). Nematodes use their stylets to secrete a variety of molecules (effectors) that suppress the defense responses in infected plant cells, leading to the formation of a functional feeding site (Juvale & Baum, 2018). Our transcriptome analysis showed that several plant genes that are activated upon infection by a variety of pathogens are expressed more abundantly in MAS as compared with FAS. Particularly relevant is a set of genes with roles in plant basal defence against pathogens. Among them are members of the polygalacturonase-inhibiting protein gene family (*PGIP1* and *PGIP2*), which are involved in perception and activation of damage-associated defense responses. Both *PGIP1* and *PGIP2* were more strongly expressed in MAS than in FAS, and loss-of-function *pgip1* mutants showed a significant increase in the average number of females and a corresponding decrease in the average number of males as compared to Col-0 (Shah et al. 2017,). Likewise, several host secondary metabolism genes, such as phytoalexin deficient (*PAD3*), indole glucosinolate omethyltransferase 1 (*IGMT1*), and cytochrome P450 91A1 (*CYP81D1*), were significantly upregulated in MAS compared to FAS. Intriguingly, we also found that PGIP-mediated changes in host susceptibility to cyst nematodes involve the activation of genes encoding enzymes for host secondary metabolism (Shah et al., 2017).

Our results suggest that ESD in cyst nematodes is influenced by the extent to which host defence responses are avoided/suppressed. We propose that some of the nematodes may be able to effectively suppress or avoid the defence response during the initial stages of infection, leading to the conditions that favor the development of females. In cases where defence responses were established by the plant, the development of males was induced. An alternative theory is that the sex of the beet cyst nematode is genetically pre-determined and male nematodes are not able to effectively suppress the defence responses below a minimum level that is required for the development of females. However, our observations showing that knocking out key defence response genes, such as *PGIP1*, increased the number of females and decreased the number of males makes it unlikely that the sex of the beet cyst nematode is pre-determined.

### MAS shows upregulation of genes involved in nutritional stress response

We found that a number of genes induced by sulfur deficiency, iron deficiency, or starvation were upregulated in MAS as compared with FAS. These observations raise the question of whether the availability of certain essential elements influences the sex differentiation of the nematodes. We propose that nematodes associated with syncytia that are unable to provide them with optimal nutrients may develop as males. This hypothesis is supported by earlier studies suggesting that a deficiency of essential elements, like phosphorous, nitrogen, and potassium, significantly increased the number of males on the host plant (Kämpfe & Kerstan, 1964). Alternatively, it is possible that nematode juveniles provoke a local host defence response during migration and establishment of the ISC and an inability to effectively suppress or overcome these defence responses may lead to restrictions in the supply of nutrient, leading to the development of males. More work will be needed to investigate this hypothesis.

### Functional characterization of candidate genes

#### IRREGULAR XYLEM 12

Our transcriptome data showed that *IRREGULAR XYLEM 12* (*IRX12*) is one of the most strongly upregulated genes in FAS compared to MAS. *IRX12* is strongly expressed in vascular bundles and is involved in constitutive lignification of the *Arabidopsis* stem by regulating phenylpropanoid metabolism (Brown et al., 2005; Berthet et al., 2011). The knock-out mutant for *IRX12* exhibits irregular xylem morphology due to the negative pressure produced by water transportation (Brown et al., 2005). The phenotype arises due to defective cellulose and lignin biosynthesis, which would otherwise provide resistance against compressive forces. The phenotype is also an indicator of secondary cell wall malformation (Turner & Somerville, 1997; Jones, Ennos, & Turner, 2001). The intensity of the phenotype of irregular or distorted xylem vessels in the *irx12* mutant can be mild to severe within the vascular bundles of the same plant (Brown et al., 2005). Moreover, *IRX12* also provides mechanical strength to xylem vessels (Yokoyama & Nishitani, 2006). Our infection assays on the knock-out mutant, *irx12*, showed a reduction in the number of females, whereas the number of males was increased significantly compared to the wild type. Because *irx12* mutants are known for defects in lignin biosynthesis and mild reductions in cellulose during secondary cell wall formation of xylem (Yokoyama & Nishitani, 2006; Hao & Mohnen, 2014), we suggest that upregulation of *IRX12* in FAS is required for increased thickening of the syncytial cell wall, which would in turn support an increased tolerance of the syncytia to high turgor pressure. In the absence of *IRX12*, an impairment of cell wall thickening may restrict the enlargement of syncytia, leading to the development of more males.

Alternatively, impairment in secondary cell wall biosynthesis might lead to the activation of plant defense pathways that restrict further development of female nematodes and favor the development of more males. This hypothesis is supported by previous studies where knocking out cell wall biosynthesis genes, such as *IRX5* and *IRX3*, conferred enhanced resistance against necrotrophic and vascular pathogens (Ellis, Karafyllidis, Wasternack, & Turner, 2002; Ramos et al., 2013).

#### GLYCOSYLPHOSPHATIDYLINOSITOL-ANCHORED LIPID TRANSFER PROTEIN 6 (LTPG6)

Non-specific lipid transfer proteins (nsLTPs) are plant-specific proteins encoded by large gene family. One of the major types are LTPGs, where the transcript encodes an additional C-terminal signal sequence, which leads to the addition of a glycosylphosphatidylinositol (GPI)-anchor via post-translational modification (Debono et al., 2009). Although the *in vivo* role of these proteins has not yet been determined, they have been shown to bind and transport lipid molecules *in vitro* (Carvalho & Gomes, 2007). Therefore, it is assumed that LTPGs are involved in lipid transport to the cell surface *in vivo* (Edstam & Edqvist, 2014). Several members of LTPGs, including LTPG6, have also been found in phloem exudates of *Arabidopsis*, secreted upon inoculation with *Pseudomonas syringae* pv. tomato. These observations led to the idea that *LTPGs* may be involved in systemic acquired resistance; however, no such role for *LTPGs* has been established (Carella et al., 2016). Nevertheless, knock-out mutants for some *LTPGs* in *Arabidopsis* have reduced fertility due to infertile ovules, an inability to restrict the uptake of tetrazolium salt, and decreased levels of ω-hydroxy-hydroxy fatty acids in the seed coat. These studies suggested that *LTPGs* may be involved in the biosynthesis or deposition of suberin or sporopollenin (Edstam & Edqvist, 2014).

Our pathogenicity assays with *ltpg6* mutants showed that although the number of female nematodes was significantly decreased, there was no significant increase in the number of male nematodes. Moreover, the size of FAS was also significantly decreased in the *ltpg6* mutants as compared to that in Col-0. Although, it is hard to identify a role for *LTPG6* in plant-nematode interactions based on current data, *LTPG6* may be involved in suberin biosynthesis and deposition around the feeding site.

#### LONGIFOLIA 1 (LNG1)

The infection assays on the *lng1* mutant revealed that the number of female nematodes was significantly reduced and the number of male nematodes was significantly increased. Moreover, the sizes of syncytia were also significantly reduced in the *lng1* mutant as compared to wild-type *Arabidopsis*. The *LNG* gene family consists of only two genes, *LNG1* and *LNG2*, in *Arabidopsis*. Indigenously, they are expressed in various plant organs, including leaves, flowers, and lateral roots where the cellular expression is localized in the cytosol and nucleus. The *LNG1-*overexpression plants have extremely long leaves, elongated floral organs, and elongated siliques. Moreover, *LNG1* and *LNG2* regulate leaf morphology by promoting longitudinal polar cell elongation (Lee et al., 2006). Kerstan (1969) studied the development of males and females in *H. schachtii* with respect to root and giant cell (syncytium) diameter and found that a certain minimum size of syncytium is required for the development of females. Based on our results, we suggest that in *lng* knock-out mutants, the diameter of the syncytium is reduced which may be less supportive to females, leading to the development of more males. Further studies involving a *lng1 lng2* double mutant may provide more insights into the role of *LNGs* in syncytium and nematode development.

## Conclusion

In the present study, we explored the host factors driving the sexual differentiation of the cyst nematode *H. schachtii* at the molecular level. We conclude that a number of factors, including the intensity of the host immune responses, availability of nutrients, and selection of ISC, may contribute to the development of females or males. In the future, it will be important to extend candidate screening to identify additional host mutants with a strong influence on the sex ratio. It will also be critical to further investigate the mechanisms by which these host genes influence the sex of other cyst nematode species. These proposed studies may provide additional resources for the development of nematode-resistant cultivars.

## Acknowledgments

The authors are thankful to Keith Lindsey and Jennifer Topping for help with LCM. We also acknowledge the excellent contribution of Stefan Neumann, Gisela Sichtermann and Thomas Gerhardt for help maintaining nematode culture.

## References

Anjam, M.S., Ludwig, Y., Hochholdinger, F., Miyaura, C., Inada, M., Siddique, S. & Grundler, F.M.W. (2016) An improved procedure for isolation of high-quality RNA from nematode-infected *Arabidopsis* roots through laser capture microdissection. Plant Methods 12, 1–9.

Anwer, M.A., Anjam, M.S, Shah, J.S., Hasan, M.S., Naz, A.A., Grundler, F.M.W., & Siddique, S. (2018). Genome-wide association study uncovers a novel QTL allele of *AtS40-3* that affects the sex ratio of cyst nematodes in Arabidopsis. Journal of Experimental Botany doi: 10.1093/jxb/ery019.

Berthet, S., Demont-Caulet, N., Pollet, B., Bidzinski, P., Cézard, L., Le Bris, P., …, Jouanin, L. (2011) Disruption of Laccase4 and 17 results in tissue-specific alterations to lignification of *Arabidopsis thaliana* stems. Plant Cell 23, 1124–37.

Böckenhoff, A. & Grundler, F.M.W. (1994) Studies on the nutrient uptake by the beet cyst nematode *Heterodera schachtii* by in situ microinjection of fluorescent probes into the feeding structures in *Arabidopsis thaliana*. Parasitology 109, 249–255.

Brown, D.M., Zeef, L.A. H., Ellis, J., Goodacre, R. & Turner, S.R. (2005) Identification of novel genes in *Arabidopsis* involved in secondary cell wall formation using expression profiling and reverse genetics. Plant Cell 17, 2281–2295.

Carvalho, A.de O. & Gomes, V.M. (2007) Role of plant lipid transfer proteins in plant cell physiology - A concise review. Peptides 28, 1144–1153.

Carella, P., Merl-Pham, J., Wilson, D.C., Dey, S., Hauck, S. M., Vlot, C., & Cameron, R. K. (2016). Comparative proteomics analysis of Arabidopsis phloem exudates collected during the induction of systemic acquired resistance. Plant Physiology, pp-00269.

Debono, A., Yeats, T.H., Rose, J.K. C., Bird, D., Jetter, R., Kunst, L. & Samuels, L. (2009) *Arabidopsis* LTPG is a glycosylphosphatidylinositol-anchored lipid transfer protein required for export of lipids to the plant surface. Plant Cell 21, 1230–8.

Den Ouden, H. (1960) A note on parthenogenesis and sex determination in *Heterodera rostoschiensis* Woll. Nematologica 5, 215–216.

Edstam, M.M. & Edqvist, J. (2014) Involvement of GPI-anchored lipid transfer proteins in the development of seed coats and pollen in *Arabidopsis thaliana*. Physiol. Plant. 152, 32–42.

Ellenby, C. (1954) Environmental determination of the sex ratio of a plant parasitic nematode. Nature 174, 1016–1017.

Ellis, C., Karafyllidis, I., Wasternack, C. & Turner, J. G. (2002) The *Arabidopsis* mutant *cev1* links cell wall signaling to jasmonate and ethylene responses. Plant Cell 14, 1557–1566.

Endo, B. (1965) Histological responses of resistant and susceptible soybean varieties, and backcross progeny to entry and development of *Heterodera glycines*. Phytopathology 55, 375–381.

Golinowski, W., Grundler, F.M.W. & Sobczak, M. (1996) Changes in the structure of *Arabidopsis thaliana* during female development of the plant-parasitic nematode *Heterodera schachtii*. Protoplasma 194, 103–116.

Grundler, F.M.W., Betka, M., & Wyss, U. (1991) Influence of changes in the nurse cell system (syncytium) on sex determination and development of the cyst nematode *Heterodera-schachtii* - total amounts of proteins and amino-acids. Phytopathology 81, 70–74.

Hao, Z. & Mohnen, D. (2014) A review of xylan and lignin biosynthesis: Foundation for studying *Arabidopsis* irregular xylem mutants with pleiotropic phenotypes. Critical Review in Biochemistry Molecular Biolology 43, 212–241.

Juvale P.S, Baum T.J (2018) “Cyst-ained” research into Heterodera parasitism. PLoS Pathogens 14: e1006791.

Hofmann, J., Szakasits, D., Blöchl, A., Sobczak, M., Daxböck-Horvath, S., Golinowski, W., …, Grundler, F.M.W. (2007) Starch serves as carbohydrate storage in nematode-induced syncytia. Plant Physiology 146, 228–235.

Holbein, J., Grundler, F.M.W. & Siddique, S. (2016) Plant basal resistance to nematodes: an update. Journal of Experimental Botany 67, 2049–61.

Hütten, M., Geukes, M., Misas-Villamil, J.C., van der Hoorn, R.A., Grundler, F.M.W., & Siddique, S. (2015) Activity profiling reveals changes in the diversity and activity of proteins in Arabidopsis roots in response to nematode infection. Plant Physiology and Biochemistry 97, 36–43.

Ithal, N., Recknor, J., Nettleton, D., Maier, T., Baum, T.J. & Mitchum, M.G. (2007) Developmental transcript profiling of cyst nematode feeding cells in soybean roots. Molecular Plant Microbe Interaction 20, 510–525.

Jones, L., Ennos, A.R. & Turner, S. R. (2001) Cloning and characterization of irregular xylem4 (IRX4): a severely lignin-deficient mutant of *Arabidopsis*. Plant Journal 26, 205–216.

Kämpfe, L. & Kerstan, U. (1964) Die Beeinflussung Des Geschlechtsverhältnisses in Der Gattung *Heterodera* Schmidt. Nematologica 10, 388–398.

Kerstan, U. (1969) Die Beeinflussung Des Geschlechterverhältnisses in Der Gattung *Heterodera*. Nematologica 15, 210–228.

Lee, Y.K., Kim, G.T., Kim, I.J., Park, J., Kwak, S.S., Choi, G. & Chung, W.I. (2006)Longifolia1 and Longifolia2, two homologous genes, regulate longitudinal cell elongation in Arabidopsis. Development 133, 4305–4314.

Mendy, B., Wang’ombe, M.W., Radakovic, Z.S, Holbein, J., Ilyas, M., Chopra, D., …, Siddique, S. (2017) Arabidopsis leucine-rich repeat receptor–like kinase NILR1 is required for induction of innate immunity to parasitic nematodes. PLoS Pathogens 13(4): e1006284.

Molz, E. (1920) Versuche zur Ermittlung des Einflusses aüsserer Faktoren auf des Geschlechtsverhältnis des Rübennematoden (*Heterodera schachtii* A. Schmidt). Landwirthschaftliche Jahrbücher. 54, 769–91.

Müller, J., Rehbock, K. & Wyss, U. (1981) Growth of *Heterodera schachtii* with remarks on amounts of food consumed. Rev. Nematologica 4, 227–234.

Offler, C.E., McCurdy, D.W., Patrick, J.W. & Talbot, M.J. (2003) Transfer cells: Cells specialized for a special purpose. Annual Review Plant Biology 54, 431–454.

Pfaffl, M.W. (2001) A new mathematical model for relative quantification in real-time RT-PCR. Nucleic Acids Research 29:e45.

Puthoff, D.P., Nettleton, D., Rodermel, S.R. & Baum, T.J. (2003) *Arabidopsis* gene expression changes during cyst nematode parasitism revealed by statistical analyses of microarray expression profiles. Plant Journal 33, 911–921.

Ramos, B., González-Melendi, P., Sánchez-Vallet, A., Sánchez-Rodríguez, C., López, G. & Molina, A. (2013) Functional genomics tools to decipher the pathogenicity mechanisms of the necrotrophic fungus *Plectosphaerella cucumerina* in *Arabidopsis thaliana*. Molecular Plant Pathology 14, 44–57.

Raski, D.J. (1950) The life history and morphology of the sugar beet nematode, Heterodera schachtii Schmidt. Phyatopathology 40, 135–152.

Sengbusch, R. (1927) Beitrag zur Biologie des Rübennematoden *Heterodera schachtii*. Zeitschrift für Pflanzenkrankheiten und Pflanzenschutz 37: 86–102.

Shah, S.J., Anjam, M.S., Mendy, B., Anwer, M.A., Habash, S.S., Lozano-Torres, J.L., …, Siddique, S. (2017). Damage-associated responses of the host contribute to defence against cyst nematodes but not root-knot nematodes. Journal of Experimental Botany 68: 5949–5960.

Siddique, S., Endres, S., Sobczak, M., Radakovic, Z.S., Fragner, L., Grundler, F.M.W., …, Bohlmann, H. (2014) Myo-inositol oxygenase is important for the removal of excess myo-inositol from syncytia induced by Heterodera schachtii in Arabidopsis roots. New Phytologist 201, 476–485.

Siddique, S., Sobczak, M., Tenhaken, R., Grundler, F.M.W. & Bohlmann, H. (2012) Cell wall ingrowths in nematode induced syncytia require UGD2 and UGD3. PLoS One 7, e41515.

Sijmons, P. C., Grundler, F.M.W., von Mende, N., Burrows, P.R. & Wyss, U. (1991) *Arabidopsis thaliana* as a new model host for plant-parasitic nematodes. Plant Journal 1, 245–254.

Sobczak, M., Golinowski, W. & Grundler, F.M.W. (1997) Changes in the structure of *Arabidopsis thaliana* roots induced during development of males of the plant parasitic nematode *Heterodera schachtii*. European Journal Plant Pathology 103, 113–124.

Sobczak M., Golinowski W, Grundler F.M.W. (1999) Ultrastructure of feeding plugs and feeding tubes formed by *Heterodera schachtii*. Nematology 1 363–374.

Szakasits, D., Heinen, P., Wieczorek, K., Hofmann, J., Wagner, F., Kreil, D.P., Sykacek, P., Grundler, F.M.W. & Bohlmann, H. (2009) The transcriptome of syncytia induced by the cyst nematode *Heterodera schachtii* in *Arabidopsis* roots. Plant Journal 57, 771–784.

Trudgill, D. L. (1991) Resistance to and tolerance of plant parasitic nematodes in plants. Annual Review Phytopathology 29, 167–192.

Trudgill, D.L. (1967) The effect of environment on sex determination in *Heterodera rostochiensis*. Nematologica 13, 263–72.

Turner, S.R. & Somerville, C.R. (1997) Collapsed xylem phenotype of Arabidopsis identifies mutants deficient in cellulose deposition in the secondary cell wall. Plant Cell 9, 689–701.

Wyss, U. (1992) Observations on the feeding behaviour of *Heterodera schachtii* throughout development, including events during moulting. Fundamental and Applied Nematology 15, 75–89.

Yokoyama, R. & Nishitani, K. (2006) Identification and characterization of *Arabidopsis thaliana* genes involved in xylem secondary cell walls. Journal of Plant Research 119, 189–194.

Yi, X., Du, Z. & Su, Z. (2013) PlantGSEA: a gene set enrichment analysis toolkit for plant community. Nucleic acids research. 41(W1):W98–W103.

Zimmermann, P., Bleuler, S., Laule, O., Martin, F., Ivanov, N.V., Campanoni, P., …, Gruissem, W. (2014) Expression Data - A public resource of high quality curated datasets representing gene expression across anatomy, development and experimental conditions. BioData Mining 7:18.

